# Input data when using neural networks to estimate lower-body torques from wearable sensors during gait: Is it of great influence?

**DOI:** 10.64898/2026.05.05.722877

**Authors:** Simon Ozan, Laetitia Fradet

## Abstract

Recent advancements in wearable sensors and machine learning show promise for estimating lower-body joint torques outside of laboratory settings. Inertial Measurement Units combined with Convolutional Neural Networks have proven effective for this task. However, the impact of different input data types and formats remains underexplored. This study investigates how variations in input data influence the prediction of lower-body joint torques during walking. Results indicate that while dataset choice causes only minor differences in prediction performance, the overall quality of the dataset plays a more critical role than the specific input variables in achieving accurate torque predictions using wearable sensors.

## 1. INTRODUCTION

Analysing joint torques in the musculoskeletal system during gait is crucial for understanding the mechanisms of human movement and identifying locomotion dysfunctions. To calculate joint torques in the lower limbs, the commonly used bottom-up inverse dynamics method involves solving the Newton-Euler equations for each segment, starting from the distal segment where the external force is applied. This method requires the measurement of kinematics and Ground Reaction Forces (GRF), typically carried out in a laboratory setting using optoelectronic systems and force plates, respectively. While these reference measurement devices offer high accuracy, they are also cumbersome, making them unsuitable for in-situ assessments.

In recent years, machine learning (ML) has emerged as a viable alternative for predicting biomechanical parameters related to human movement—such as joint torques—using data collected from wearable sensors, opening up opportunities for in-situ assessments. Various ML algorithms have been documented in the literature for predicting lower-limb joint torques, such as Artificial Neural Networks [1–5], Multilayer Perceptron [6], Recurrent Neural Network [6], Long Short-Term Memory [3, 6, 7], Support Vector Machine [8], Random Forest [8], Multivariate Adaptive Regression Splines [8], Temporal Convolutional Networks [3, 7], a specific deep learning method [9], and Convolutional Neural Networks (CNNs) [3, 6–8, 10]. To determine which type of ML algorithm is most effective for predicting joint torques from wearable sensor data, several studies have proposed training and comparing different ML algorithms [3, 6–8]. In these studies, CNNs consistently rank among the most effective—if not the most effective—ML algorithms for predicting lower-body joint torques.

While previous studies provide insights into selecting appropriate ML methods, another unresolved issue concerns the choice of input data. ML-based approaches typically involve two steps in which input data play a crucial role. The first step is training the model using measured input and output data. In the second step, the trained model is used to predict output data from newly measured input data. In the literature, input data have been obtained from various wearable sensors—sometimes combined—such as pressure insoles used to predict GRF and subsequently calculating joint torques [11], IMUs [1, 6–8, 10, 12], goniometers [7, 12], electromyography [8, 12, 13], and even electroencephalograms [13, 14]. Due to their recent advancements and portability, IMUs have seen widespread application, as confirmed by a review [15].

Currently, even with the selection of IMUs as sensors, there are still questions regarding the choice and handling of input data. The quantity of IMUs and their placement varied significantly from one study to another. For example, two studies used a single IMU positioned on the lower-back [4, 5], but joint torques were only estimated during the stance phase. Another study employed between one and three IMUs to assess the effect of varying configurations [3]. Additionally, one study implemented three IMUs situated at the pelvis and on both feet [16]. Some investigations incorporated five IMUs positioned on the pelvis, both thighs, and shanks [6, 17], while others added extra IMUs on the feet [10]. In the works of Moghadam et al. [18] and Liang et al. [3], the impact of the number of IMUs on the prediction of joint torques during gait was assessed. The findings revealed that using all available IMUs did not enhance the precision of joint torques prediction.

Various input data measured from IMUs have also been proposed in the literature, such as the three components of linear acceleration and angular velocity expressed in the IMUs’ reference frames [1, 6], sometimes combined with the norm or mean of linear acceleration and angular velocity [3]. One study also proposed expressing acceleration and angular velocity in the underlying segment’s reference frame to describe movement in the sagittal plane [10]. This approach reduces the impact of variations in sensor placement and enables the extraction of data within the plane where the greatest movement amplitude occurs. However, it requires additional calibration movements to align sensor data with the corresponding body segment.

To the best of our knowledge, while feature extraction methods have been proposed to identify the most pertinent features for use in ML [8, 18], it remains unclear whether all IMU data should be used as inputs. Furthermore, the influence of the reference frame in which the data is expressed has not been evaluated.

More is not always better, and the impact of data processing and the quantity of input data remains uncertain. Therefore, the objective of this study was to build upon the results of Moghadam et al. [18] and Liang et al. [3] to determine whether the expression, number, and type of input IMU data influence the prediction of lower-body joint torques during locomotion. This study also proposes evaluating the potential contribution of pressure insole data to the input dataset, as insole pressure provides partial information about ground reaction forces and therefore contains information on kinetics, not just kinematics.

Since convolutional neural networks (CNNs) appear to be particularly effective in predicting joint torques, a CNN was selected as the machine learning algorithm for this study

## 1 METHODS

### 1.1 Participants

Nineteen volunteers (10 female and 9 male) participated in the study (age: 47.5 ± 22.1 years; height: 169.2 ± 9.7 cm, weight: 63.7 ± 10 kg). The participants varied in age, physical characteristics, and physical ability to avoid drawing conclusions that are too specific to a particular population.

All participants provided informed consent prior to participating in the study. The study was approved by an institutional review board (CER-TP 2022-01-07).

### 1.2 Data acquisition

Participants wore two pressure insoles in their shoes (250 Hz, Ultium Insoles, Noraxon) and had 7 IMUs (128 Hz, APDM) placed on their pelvis, thighs, shanks, and feet, as illustrated in Fig. 1. The IMUs were visually aligned with the respective underlying segments. Each participant was also outfitted with 36 reflective markers, following the Conventional Gait Model version 2.1, to capture body segment motion using 19 infrared cameras (100 Hz; T170, Vicon, Oxford Metrics).

**Figure 1.**
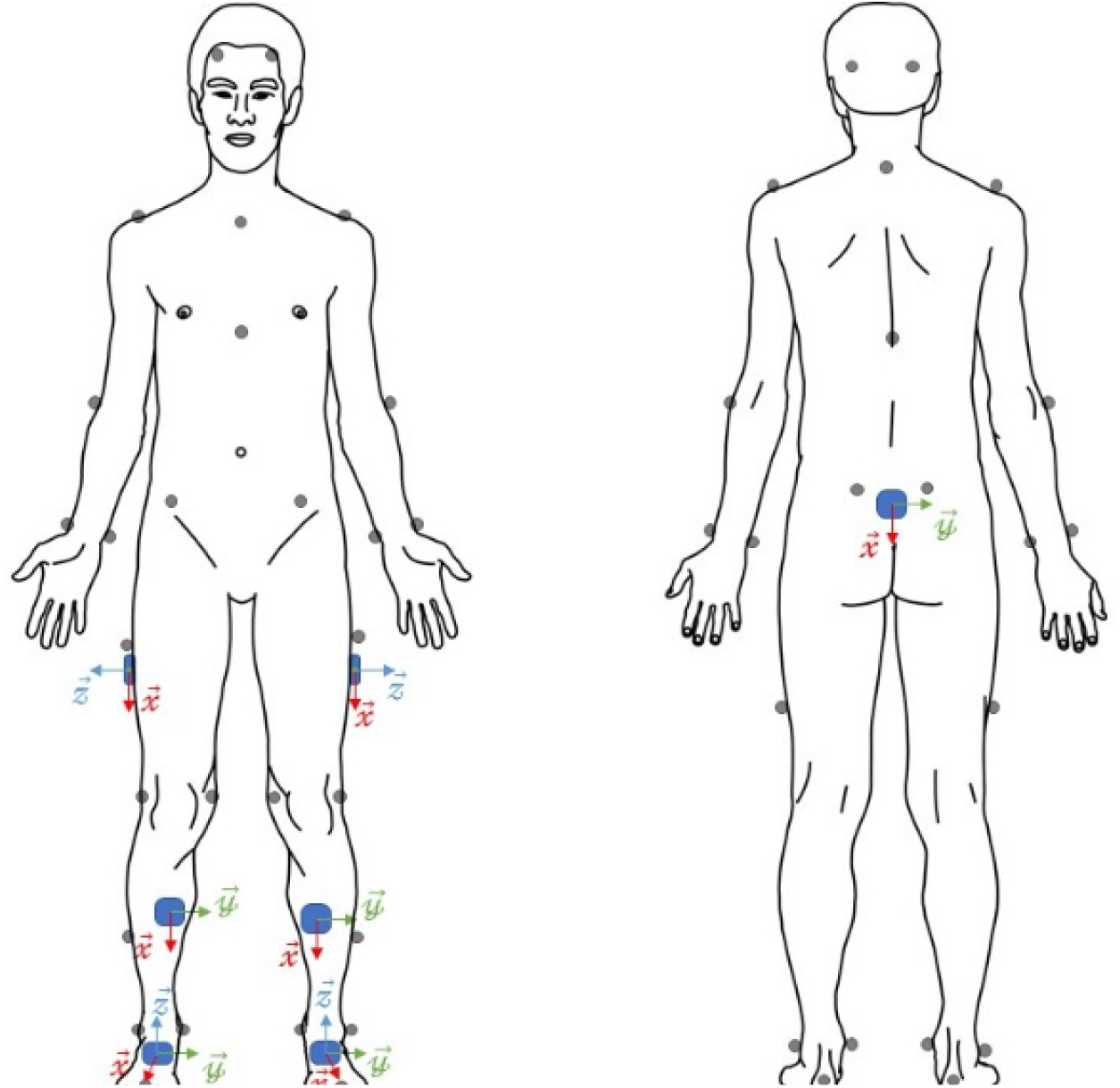
Reflective markers and IMU placement for data collection.

They had to walk across 10 force plates (Sensix, Poitiers) to measure Ground Reaction Force (GRF) for each footstep.

Each participant completed 10 walking trials under various instructions: walking at a self-selected speed, at a slow pace, and at a rapid pace, with their feet positioned either normally, wider, or narrower than usual.

### 1.3 Gold Standard method

Marker trajectories and GRF data were filtered using a second-order zero-lag Butterworth low-pass filter with a cut-off frequency of 15 Hz [19]. Subsequently, hip flexion/extension, knee flexion/extension, abduction/adduction, and ankle dorsi/plantarflexion torques were calculated using the Conventional Gait Model [20] and normalised to the subject’s body mass.

The data were segmented into gait cycles based on foot strikes identified from vertical force data, using a threshold of 15□N. Computed joint torques were time-normalized to 101 points per cycle via linear interpolation. For each subject and condition, data from six right-side gait cycles were retained.

### 1.4 Machine Learning method

#### 1) Data Processing of the wearable sensor data

As will be further explained, two representations of the inertial data were used. In one case, the data were expressed in the IMU reference frame (corresponding thus to the raw data); in the other, they were expressed in the anatomical segment reference frame. For the latter, the sensor-to-segment calibration method used was the “Postures” method, which involved recording the subject in two specific postures: a standing posture, during which the subject stood with feet parallel, and a sitting posture, during which the subject sat with legs extended [21].

Wearable sensor data were segmented into gait cycles, aligned with those defined by the optoelectronic system, using foot strikes identified from insole pressure data. The data were then time-normalized to 101 samples via linear interpolation.

#### 2) Convolutional Neural Network

A CNN is a type of neural network characterized by multiple layers that processes structured data arrays. Distinct CNNs were developed for each array generation method and for each output variable namely, the hip and knee flexion/extension torques, the ankle dorsi/plantarflexion torque, and the knee abduction/adduction torque. All the CNNs were initialised before training with the same architecture and hyperparameters as a CNN proposed in the literature [10]. The CNNs were subsequently trained to predict output variables from the arrays generated using the datasets and methods outlined below. Training of the CNNs was performed using data from 80% of the subjects (i.e., 15 subjects), resulting in a total of 450 gait cycles. A four-fold cross-validation approach was used to train the CNNs.

#### 3) Inputs datasets and arrays

Six different datasets were considered in the study to generate the structured data arrays used as inputs of the CNNs.

The first dataset (DS1) corresponds to the dataset and method presented in a previous study [10], which considers only inertial data in the segments’ sagittal plane. It includes then acceleration along the longitudinal and anteroposterior axes of the segment’s reference frame, as well as angular velocity around the mediolateral axis. Thus, three features are used for each of the four IMUs considered -namely, the pelvis, right thigh, right shank, and right foot (Table 1).

**Table 1:**
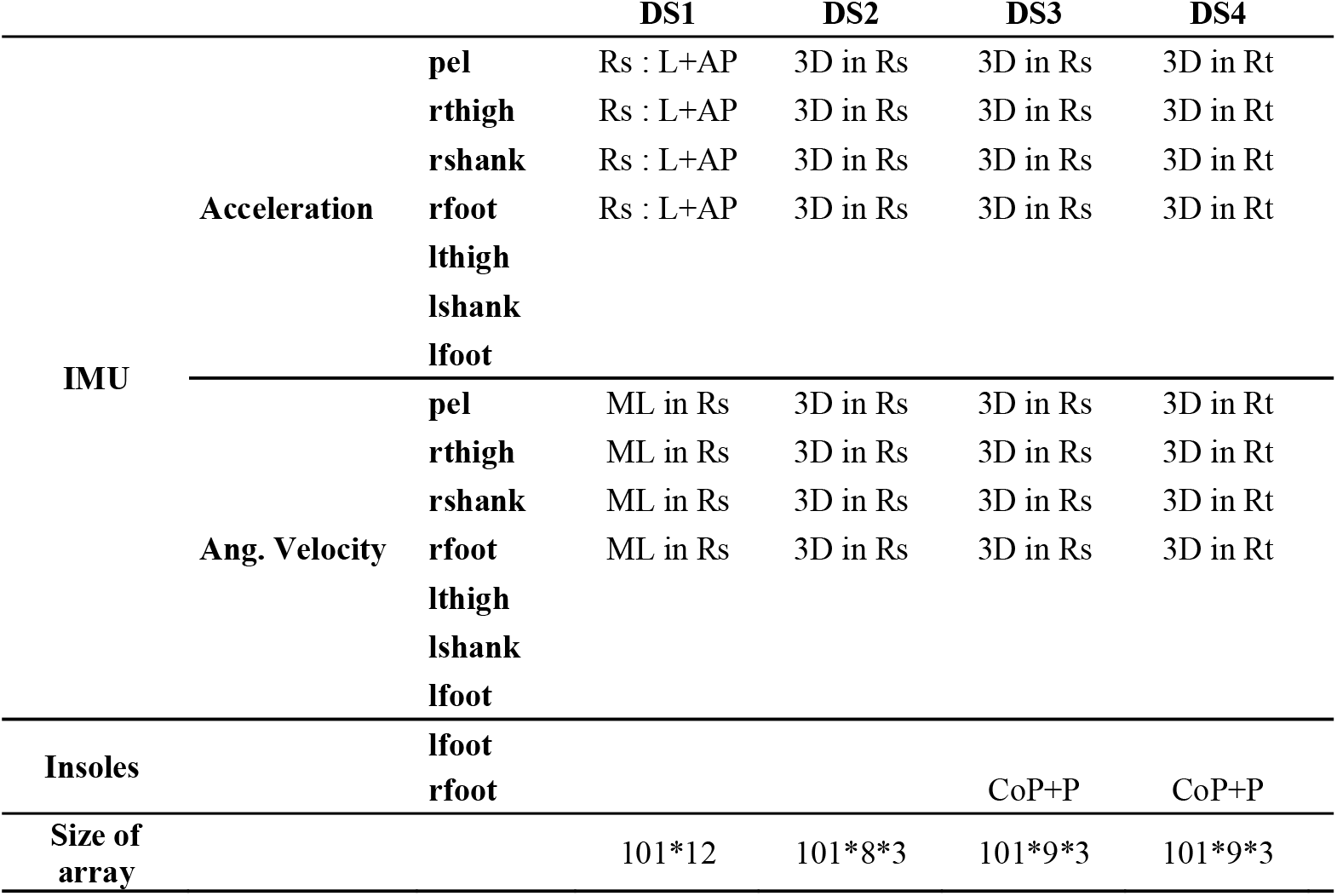
Dataset and image creation for each one of the methods compared. *Rs* signifies that the acceleration or the angular velocity were expressed in the underlying segment frame, and *Rt* that they were expressed in the sensor technical frame. *AP, ML*, and *L* stand for the segment antero-posterior, medio-lateral, and longitudinal and axes respectively. P represents the insole pressure. *Pel, rthigh, rshank, rfoot, lthigh, lshank, lfoot* stand for the pelvis, right thigh, right shank, right foot, left thigh, left shank, and left foot respectively.

The data were normalized using the min-max normalization method (1).

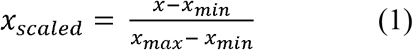

The input data therefore consisted of a 101 (frames) × 12 (IMU features) 2D array, which is equivalent to a grayscale image.

For the second dataset (DS2), the full 3D accelerations and angular velocities expressed in the underlying body-segment reference frame were used. After applying the same min-max normalization procedure, the 3D data were structured as a 101 (frames) × 8 (IMU locations) × 3 (axes) 3D array, which is analogous to RGB image data [6].

For dataset 3 (DS3), insole data were integrated with the IMU data from DS2. The mediolateral (Xcop) and anteroposterior (Ycop) positions of the Centre of Pressure (CoP) were normalized based on foot width and length, respectively. The final dimension of the insole data represented the total pressure recorded by the insole, normalized according to the pressure range. These three channels underwent normalization as previously outlined and were incorporated into the matrices of DS2, yielding a 3D array of dimensions 101 (frames) * 9 (data) * 3.

For dataset 4 (DS4), the same approach as DS3 was applied, with the notable difference that sensor-to-segment calibration was not executed. Consequently, the IMU data were represented in the IMU reference frame rather than the segment reference frame as in DS3.

DS5 is based on the input data used by Donahue and Hahn [16], namely the 3D accelerations and angular velocities, as well as their magnitudes (norms), but only for the IMUs located on the pelvis and both feet. The input data in this case consisted of a 101 (frames) × 24 (features) 2D array.

DS6 builds upon DS4, with the addition of data from the left foot IMU.

### 1.5 Evaluation

Each CNN was assessed using 120 gait cycles from the four remaining subjects. To evaluate the predictions made by the various CNNs, the Root Mean Square Difference (RMSD) and the Pearson Correlation Coefficient (CC) were calculated between the estimated values and the joint torque obtained through the Gold Standard method, which served as the reference. Additionally, the difference between the peak values obtained from the reference and each dataset was calculated. For the positive torque values, the RMSD reflected the differences in peak values for hip flexion, knee flexion and adduction, as well as ankle dorsiflexion torques. Conversely, for the negative values, it represented the peaks of hip extension, knee extension and abduction, along with ankle plantarflexion torques.

### 1.6 Statistical analysis

The normality of the data was assessed using the Shapiro-Wilk test, while the equality of variances was evaluated using the Bartlett test. A one-way ANOVA was conducted to examine the impact of the dataset on the predictions. The significance level was established at p<0.05. In cases of significance, a post-hoc test with a Bonferroni correction was applied to the data.

## 2 RESULTS

At the top of Fig. 2, an example of torque temporal evolution during a gait cycle from subject 17 is shown. For this subject, the torque patterns obtained from the different datasets appear similar. At the bottom of Fig. 2, boxplots summarize the descriptive statistics for each torque.

**Figure 2.**
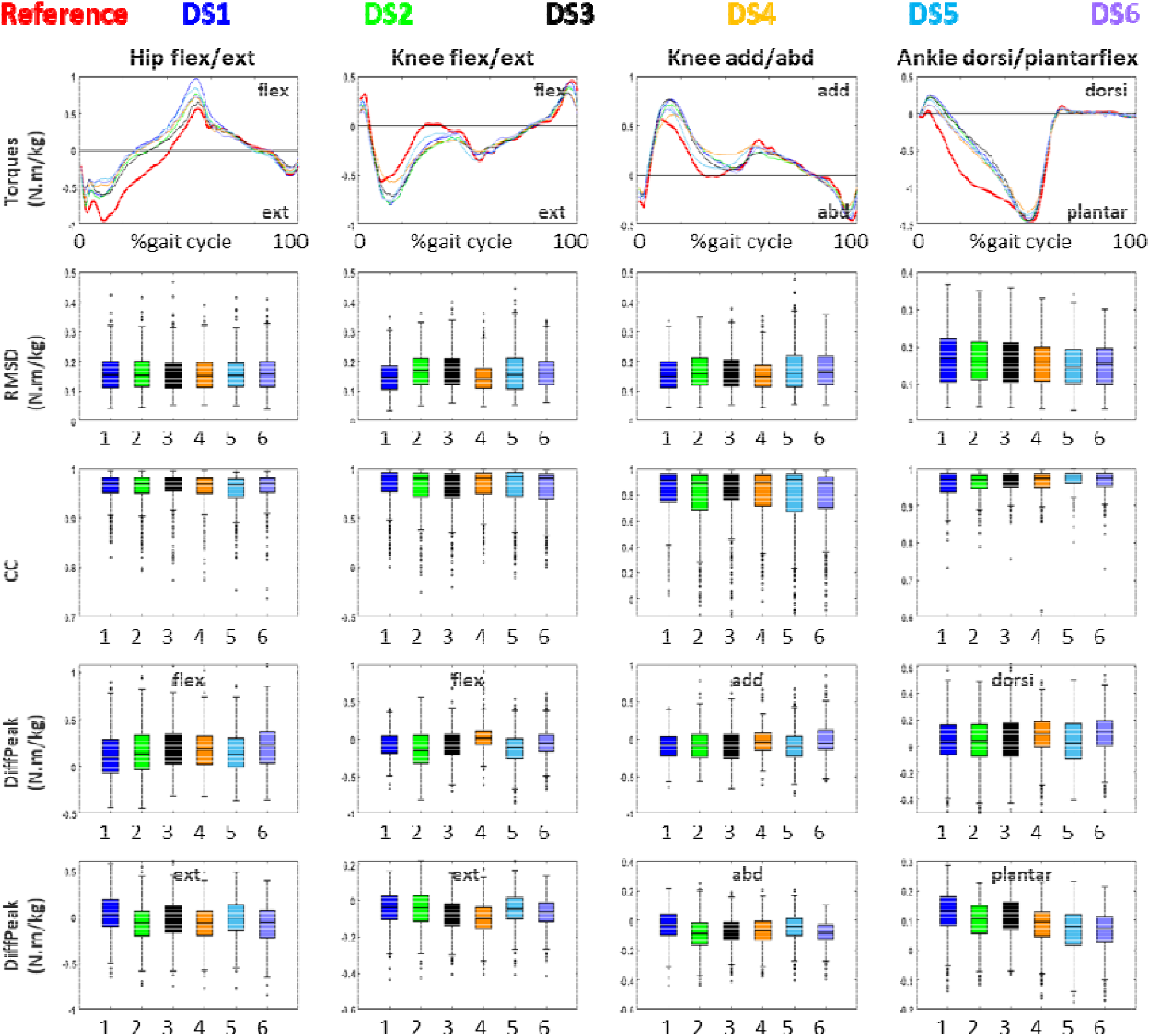
Top line: Torques obtained for a gait cycle from subject 17 with, in red, the reference, in purple, the dataset DS1, in light blue, DS2, in orange, DS3, in black, DS4, in green, DS5, and in blue, DS6. Subsequent lines: Median, 25th and 75th percentiles and outliers for the RMSD, CC, differences in peak values between each dataset, and the reference for the positive and negative values.

The mean values and standard deviation, as well as the results of the statistical tests, are presented in Table 2.

**Table 2:**
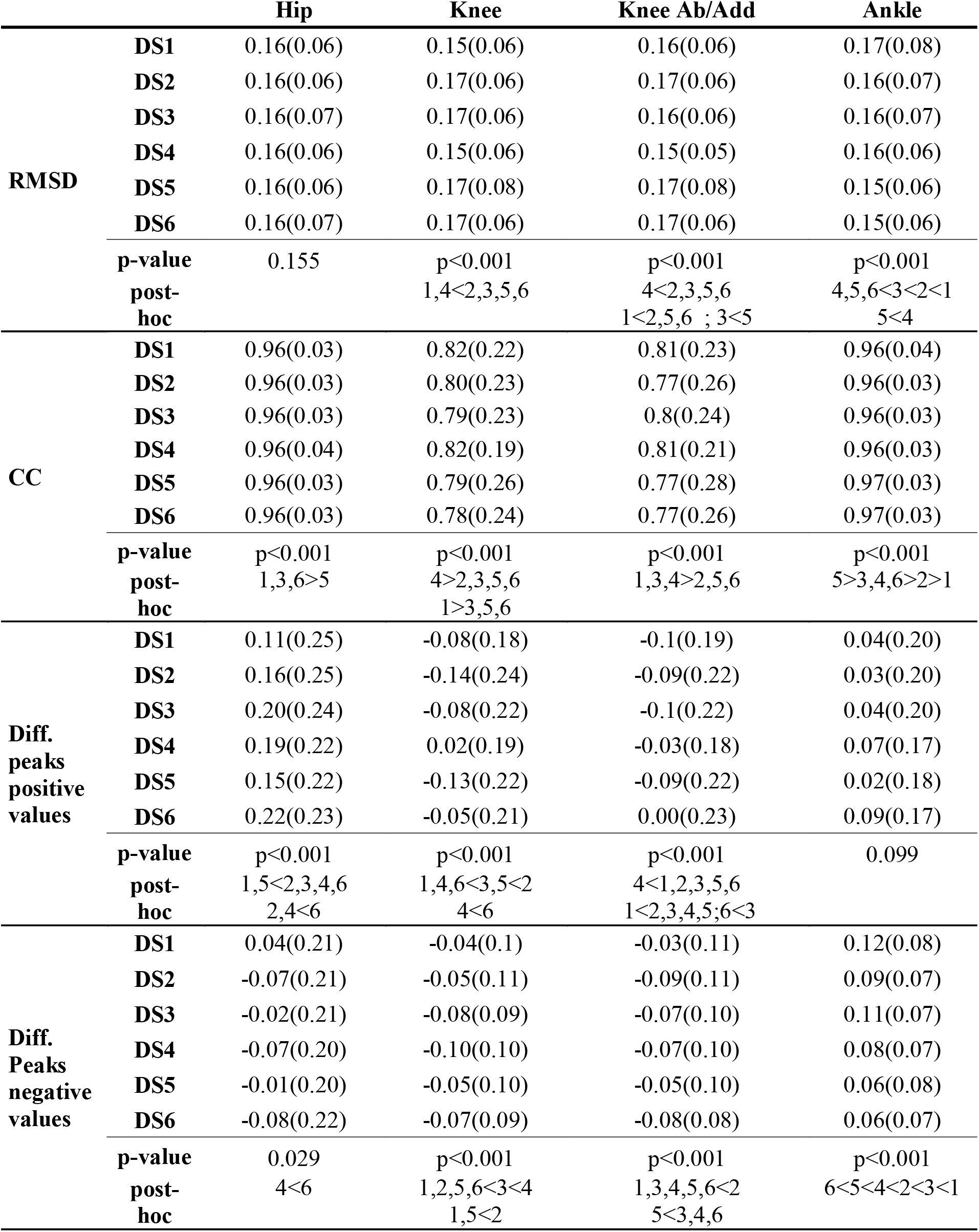
Mean values (standard deviation) and results of the statistical tests. The negative values for the peak differences indicate an underestimation relative to the reference, though the statistics were calculated using the absolute difference.

The dataset significantly impacted the RMSD results for each torque (p < 0.001), except for the hip flexion/extension torque (p = 0.155). According to the post-hoc tests, for the knee flexion/extension torque, the RMSD was smaller for DS1 and DS4 compared to the other datasets. For the knee abduction/adduction torque, DS4 exhibited a significantly smaller RMSD than all other datasets, except DS1. DS1 had a smaller RMSD than DS2, DS5, and DS6, while DS3 showed a smaller RMSD than DS5. In the case of the ankle dorsiflexion/plantarflexion torque, the RMSD was lowest for DS4, DS5, and DS6, followed by DS3, DS2, and DS1. DS5 had also a smaller RMSD than DS4.

The dataset also influenced the correlation coefficient (CC) at all joints. For hip flexion/extension, DS1, DS3, and DS6 showed larger CCs than DS5. In knee flexion/extension, DS1 and DS4 had higher CCs than DS3, DS5, and DS6, with DS4 also showing a statistically greater CC than DS2. For knee abduction/adduction, DS1, DS3, and DS4 exhibited larger CCs compared to the other dataset. Regarding ankle dorsiflexion/plantarflexion, DS5 had the highest CC, followed by DS3, DS4, and DS6, which were all greater than DS2, and DS1 showed the smallest CC.

The dataset also influenced the difference in peak torque values relative to the reference for all torques (p < 0.001), except for ankle dorsiflexion (p=0.099). For hip flexion torque, the difference from the reference was smaller for DS1 and DS5 than for all other datasets, with DS2 and DS4 showing smaller differences than DS6. In hip extension torque, DS4 had a smaller difference than DS6. For knee flexion torque, the smallest differences from the reference were observed for DS1, DS4, and DS6, compared to DS3 and DS5, which were smaller than DS2. DS4 also had a smaller difference than DS6. For knee extension torque, DS1, DS2, DS5, and DS6 had smaller differences than DS3, which in turn was smaller than DS4. Additionally, DS1 and DS5 showed smaller differences than DS2. Regarding knee abduction torque, DS4 had the smallest difference with the reference, while DS1 had a smaller difference than DS2, DS3, and DS5. DS6 demonstrated a smaller difference than DS3. For knee adduction torque, DS2 had the largest difference, while DS5 exhibited a smaller difference than DS3, DS4, and DS6. Lastly, for plantarflexion torque, DS6 had the smallest difference with the reference, followed by DS5, DS4, DS2, DS3, and DS1.

## 3 DISCUSSION

The present study proposes to test the effects of different input data expressions, numbers, and types on the prediction of lower-body kinetics during gait, using a CNN. Recent studies have used machine learning to predict lower-body kinetics during various physical activities, and showed that CNNs had among the best performances in prediction [3, 6, 18]. However, while IMUs have established themselves as preferred data sources, the impact of input data on predictions has not been fully determined. Moreover, the potential contribution of pressure insole data to the input dataset has yet to be tested, since pressure provides partial information on ground reaction forces and thus contains information on kinetics. Therefore, to complement the study by Moghadam et al. [18] and that of Liang et al. [3], which examined the effect of the number of IMUs used on lower-limb joint torque predictions, we also propose to investigate the influence of the reference frame in which the data are expressed and also the use of insole data.

According to the results, the RMSD comparing the predicted data and the original data remain high, since they are around 10% of the peak values for the hip flexion/extension and the dorsi/plantarflexion torques, and around 19% for the knee flexion/extension and abduction/adduction torques (RMSD values, expressed as a percentage of the peak values, are presented in Table A1 in the Annex). The results are quite similar to what can be seen in the Mundt et al’ paper [6] for the hip flexion/extension and ankle dorsi/plantarflexion, but seem slightly higher for the knee flexion/extension. This is also the case for the correlation coefficients, which seem worse for the present study at the knee joint compared with Mundt et al. This might be explained by the much greater training set since it was built in the Mundt et al’ study with 93 subjects, and was augmented using simulation [6]. The population used to obtain the data was also younger and slightly less heterogenous than that of the present study.

The results closely align with those presented by Dorschky et al. [10]. Notably, our study shows a slightly better RMSD for the hip flexion/extension moment, but a slightly lower CC for the knee flexion/extension moment. This discrepancy may be again attributed to the greater heterogeneity of our subject pool, which included both young and elderly participants.

It can be observed that the CCs are lower for knee flexion/extension and abduction/adduction torques. This may be due to greater time variations in these torques, which are more pronounced than those in hip flexion/extension and ankle dorsi/plantarflexion torques, as shown in Figure 2. However, we also computed the percentage of cycles for which the CC was greater than 0.9 for each torque and each dataset, as well as the percentage of cycles for which the RMSD normalised to the peak torque was lower than 10%. The results show that the predictions are really worse for the knee flexion/extension and abduction/adduction torques than for the two others. For instance, no dataset was able to predict more than 35% of gait cycles with a normalised RMSD inferior to 10% (see Table 3).

**Table 3:** Percentage of trials with a normalised RMSD less than 10% and a CC greater than 0.9 when compared to the reference for each method.

|  |  | Hip | Knee | Knee Ab/Ad | Ankle |
| --- | --- | --- | --- | --- | --- |
| **RMSD<10%** | **DS1** | 67 | 35 | 35 | 49 |
|  | **DS2** | 62 | 31 | 30 | 54 |
|  | **DS3** | 66 | 27 | 31 | 52 |
|  | **DS4** | 63 | 29 | 30 | 52 |
|  | **DS5** | 66 | 35 | 32 | 58 |
|  | **DS6** | 63 | 24 | 20 | 52 |
| **CC>0.9** | **DS1** | 95 | 57 | 55 | 93 |
|  | **DS2** | 93 | 49 | 46 | 95 |
|  | **DS3** | 96 | 48 | 51 | 96 |
|  | **DS4** | 92 | 54 | 49 | 96 |
|  | **DS5** | 92 | 55 | 54 | 96 |
|  | **DS6** | 95 | 51 | 48 | 96 |

According to the results, the predictions can be considered generally better in terms of CCs than in terms of RMSD. For example, for the hip torque, more than 96% of gait cycles predicted had a CC with the reference higher than 0.9, while only 67% resulted in a normalised RMSD lower than 10%. This might be due to the min-max normalisation method as well as the time normalization of the gait cycle, which might hinder information relative to the gait speed. Gait speed seems indeed to affect the values more than the patterns of the torques, especially at the knee [22], which could also explain why the knee torques had the worst results. This means that another proposal should be tested to normalize the data or that the data should not be normalized.

Regarding the original question, according to the statistics, the input dataset -manipulated here in terms of the number of sensors, the reference frame in which the data were expressed, and the types of data, namely inertial data possibly complemented by data from pressure insoles-does influence the prediction of lower-body torques during gait using a CNN. Our results tend also to indicate that the same dataset is not the best for all torques. Nevertheless, one can question the significance -although statistical as aforementioned-of the impact of the input dataset. Indeed, in terms of RMSD and of CC, the differences are really small between the different datasets. For instance, when looking at the means, in terms of RMSD, the differences between the methods do not exceed 0.02 N.m/kg and, in terms of correlations, the differences are inferior to 0.02 for the hip flexion/extension and ankle dorsi/plantarflexion torques and inferior to 0.04 for the knee flexion/extension and knee abduction/adduction torques.

Given the varying performance of the input datasets across the different joints and the small differences between them, drawing a definitive conclusion about the best dataset is challenging. According to our results, DS1, which corresponds to that proposed by Dorschky et al [10], which includes acceleration and angular velocity of IMUs placed at the pelvis, thigh, shank, and foot of one leg, the acceleration and angular velocity expressed in the segment sagittal plane seems the best for predicting the hip flexion/extension torque, but worst for predicting the ankle dorsiflexion/plantarflexion torque. DS4 performs best for knee flexion/extension and abduction/adduction, and DS5 for ankle dorsiflexion/plantarflexion.

At the hip, DS1 outperforms DS2 and DS3, likely due to the use of 2D data expressed in the segment reference frame for DS1, compared to the 3D data used for DS2 and DS3. This difference may stem from the fact that IMU sensor-to-segment calibration is known to yield less accurate kinematics, particularly in the frontal and transverse planes, when compared to optoelectronic systems [23]. As a result, the inclusion of less accurate 3D data in DS2 and DS3 may degrade the quality of the predictions relative to DS1.

For knee flexion/extension and abduction/adduction torques, DS4 provides slightly better predictions than the other datasets. This could be attributed to the inclusion of insole pressure data, which may indirectly offer insights into kinetics and also gait speed. As mentioned earlier, knee torques are particularly sensitive to gait speed [22]. Insole pressure data also reflect gait phases, which are influenced by gait speed specifically, the stance phase shortens as gait speed increases [24].

Regarding ankle dorsiflexion/plantarflexion torques, DS5 -based on the approach proposed by [16]-appears to provide slightly better predictions. This dataset uses only 3 IMUs, placed at the pelvis and on both feet. It can be inferred that adding data from sensors not located near the joint of interest may reduce the accuracy of predictions, as the additional data could introduce noise or less relevant information.

These observations and hypotheses lead to broader conclusions. Specifically, expressing data in the underlying segment reference frame is intended to account for slight variations in sensor positioning across subjects, and consequently, between the training and test datasets. In this study, the sensors were consistently positioned in the same location on the segments. As a result, the data seem to appear sufficiently stable such as no significant additional insights were gained from using the segment-based reference frame. Since defining the segment reference frame requires a specialized procedure, the sensor-to-segment calibration [21], one might opt not to perform this calibration.

Another key conclusion from this study is that more data is not always better. This was also observed in our study for the prediction of ankle dorsi/plantarflexion torque, which was better when using only three IMUs. Previous research has shown that using additional IMUs does not necessarily enhance predictions for kinematics [25] or the kinetics of the lower limbs [3, 18]. The reason for this may be that adding more IMUs results in redundant data, which can introduce noise or unnecessary complexity without necessarily providing meaningful additional information for prediction. However, significantly reducing the number of IMUs to just one appears to lead to suboptimal prediction performance, as reflected by the RMSD and CC values in studies by Lee and Park [3–5]. A potential resolution could be to utilise a greater number of sensors, albeit in specific combinations – potentially different combinations – for each joint torque.

To conclude on the influence of the dataset, based on the present results it cannot be excluded that some other data combination or treatment might slightly improve the prediction. However, the present results tend to suggest that the significance of the dataset on the prediction would probably remain small. Therefore, the choice of the input dataset might depend on the material available. We can just notice that if insole data do not improve the prediction of the kinetics substantially, their use is convenient to define the gait cycles.

It is also noteworthy that input data processing was minimized in the present study. As mentioned in [26], another approach involves incorporating physics knowledge into the process. In the study by Lim et al. [5], data from a single IMU located on the lower back were used to estimate the centre of mass position, velocity, and acceleration, which then served as input for the ML algorithm. Although the normalized RMSD was close to 10% with just one IMU, the correlation coefficients (CC) rarely exceeded 0.90. While promising, the study had two main drawbacks: first, during IMU data pre-processing, it made the significant assumption that gait was performed at a constant speed. This assumption was realistic in their study, as subjects walked on a treadmill, but is less applicable to real-life conditions. Second, it predicted joint torques only during the stance phase.

More generally, the value of this approach depends on the suitability and quality of the integrated physical data. For example, the model used by Lim et al. to estimate the centre of mass kinematics may not be suitable for pathological subjects.

With minimally processed input data, as in the present study, the quality of the prediction appears to depend more on the database than on the input dataset. This is illustrated by additional results presented in Annex 1. In these results (Table A2), 80% of the gait cycles were used to train the CNNs and 20% for testing, without regard to whether the data came from the same or different subjects. In this case, all results improved, with correlation coefficients (CCs) exceeding 0.92 for knee flexion/extension and abduction/adduction. This suggests that the datasets and trained CNNs were likely able to predict gait cycles from the same individuals seen during training.

As aforementioned, the prediction is probably better in the Mundt et al’ paper [6] because of the size of their training set. Previous results have shown that with CNN, the larger the training set, the better the classification performance [27]. To improve the database, attempts have been made to supplement measured data with simulated data [6, 10]. In Mundt et al. [6], a slight rotation was applied to the initial data to simulate a slight change in the sensor orientation. In Dorschky et al. [10], ‘new mechanically realistic data’ were obtained through simulation using a musculoskeletal model. In Dorschky et al., the impact of the simulated data was tested, showing a contrasting effect on joint torque prediction. Mundt et al. did not test the impact of their simulated data but since Mundt et al.’s study yielded the best predictive results in the literature, it leaves the question open regarding the value of simulated data for expanding the database. As mentioned by Dorschky et al. [10], their simulation-aided estimation of joint moments was probably limited by inaccuracies of the biomechanical model. Improving the physics-based model may further increase the benefit of simulated data. In the present study, we did not simulate data to bring variations but asked the subjects to walk in different conditions in order to induce some sort of variation in the movement. The subjects were thus asked to walk at a slower, then at a faster pace than their comfortable pace and, additionally, to walk with larger and smaller step widths. What must also be mentioned is that the training sets must be balanced [28], meaning that the same quantities of data corresponding to specific patterns must be present.

## 4 CONCLUSION

This study examined how different input data expressions, quantities, and types influence the prediction of lower-limb joint torques during walking using Convolutional Neural Networks (CNNs). While the effects of manipulating the input dataset were statistically significant, their impact on torque prediction was modest. The findings indicate that other factors, such as the quantity and balance of the training data, likely have a more substantial impact on prediction accuracy than the datasets themselves.

## Fundings and acknowledgments

The authors would like to thank the participants of this study for their contribution, and Ms Hogan for proofreading the article. This study was supported in part by the New Aquitaine region and the European Union through the Habisan program (CPER□FEDER).

## ANNEX

**Table A1:** RMSD values, expressed as a percentage of the peak values.

|  | Hip | Knee | Knee Ab/Ad | Ankle |
| --- | --- | --- | --- | --- |
| **DS1** | 9.0(3.9) | 16.4(10.8) | 17.1(11.1) | 10.5(5.0) |
| **DS2** | 9.5(4.0) | 18.1(11.5) | 18.7(12.4) | 10.3(4.5) |
| **DS3** | 9.1(3.7) | 18.6(11.6) | 17.8(11.7) | 10.0(4.5) |
| **DS4** | 9.5(4.0) | 17.0(10.1) | 17.2(10.7) | 9.9(4.2) |
| **DS5** | 9.2(3.8) | 17.7(12.1) | 18.8(12.7) | 9.4(3.9) |
| **DS6** | 9.6(3.9) | 18.2(11.0) | 19.0(11.1) | 9.9(4.1) |

**Table A2:** RMSD and CC mean values (standard deviation) when 80% of the gait cycles were taken in the training set and not the gait cycles of 80% of the subjects for the first four datasets.

|  |  | Hip | Knee | Knee Ab/Ad | Ankle |
| --- | --- | --- | --- | --- | --- |
| **RMSD** | **DS1** | 0.12(0.05) | 0.09(0.04) | 0.07(0.03) | 0.08(0.03) |
|  | **DS2** | 0.12(0.05) | 0.09(0.04) | 0.07(0.03) | 0.08(0.04) |
|  | **DS3** | 0.12(0.06) | 0.09(0.04) | 0.07(0.03) | 0.08(0.04) |
|  | **DS4** | 0.11(0.06) | 0.08(0.04) | 0.06(0.03) | 0.07(0.03) |
| **CC** | **DS1** | 0.97(0.03) | 0.95(0.06) | 0.92(0.10) | 0.99(0.01) |
|  | **DS2** | 0.96(0.03) | 0.95(0.08) | 0.92(0.11) | 0.99(0.01) |
|  | **DS3** | 0.96(0.04) | 0.95(0.07) | 0.91(0.12) | 0.99(0.01) |
|  | **DS4** | 0.97(0.03) | 0.95(0.08) | 0.93(0.09) | 0.99(0.01) |

